# CLR-Seq: a pipeline to identify bacterial microbiota species with immune-relevant glycan moieties through human C-type lectin receptor interaction

**DOI:** 10.1101/2025.03.31.646320

**Authors:** Jasper Mol, Rob van Dalen, Yvonne Pannekoek, Malgorzata E. Mnich, Marcel R. de Zoete, Mark Davids, Hilde Herrema, Nina M. van Sorge

## Abstract

Bacterial glycans are key components in immune interactions. We lack insight into the diverse glycan landscape present in complex microbial communities since current-omics techniques do not capture this post-translational information. Here we employed C-type lectin receptors (CLRs), which are dedicated innate glycan-sensing receptors, as probes for bacterial cell sorting in combination with 16S rRNA gene sequencing to identify microbiota species with a specific CLR-reactive glycan profile. We established our experimental CLR-sequencing (CLR-seq) pipeline using soluble fluorescently-labeled human macrophage galactose C-type lectin (MGL, CD301) and langerin (CD207). Both receptors identified known langerin- or MGL-interacting *Staphylococcus aureus* strains in a synthetic microbial community even when present at low abundance. Subsequent application of CLR-seq on fecal microbiota samples from healthy donors identified specific langerin- and MGL-interacting bacterial species that were subsequently validated as monocultures. In summary, CLR-seq is a modular platform that allows identification of human microbiota species based on CLR-interacting glycans with easy expansion to other CLRs or microbiota samples from patients. Given that CLRs are densely expressed on dendrites of antigen-presenting cells, this pathway may play a role in cross-barrier recognition and sampling of the environment during homeostasis.

## Introduction

The human body hosts a large variety of bacteria, collectively known as the microbiome. The composition of the microbiome is associated with health benefits but also pathologies through changes in metabolites, digestive capacity or altered inflammatory properties [1-3]. To maintain homeostasis, there is a reciprocal interaction between the microbiota and the local immune system. In addition, microbiota-triggered antibody responses even extend into the systemic compartment offering protection against pathogens [4,5]. Application of - omics technologies has allowed the identification of species and associated transcriptomes, proteins and metabolites within microbiota samples, allowing association of certain-omics signatures with specific pathologies. Further studies have subsequently been able to functionally underpin specific microbes or metabolites to health and disease processes. In contrast to this wealth of information, we currently have limited insight into the microbiota glycome, i.e. the glycan composition of the human microbiome and how this impacts human health and disease.

Glycans represent an integral part of the bacterial cell wall, providing bacterial integrity and viability [6]. Bacterial glycans are often distinct from their human counterparts and the specific glycan composition can contribute to tolerogenic or homeostatic signals through interaction with pattern recognition receptors (PRRs) on innate immune cells [7,8]. In the context of infection, it is well established that the expression of capsular polysaccharides, contributes to bacterial virulence and immune escape [9]. Consequently, directing specific antibody responses towards bacterial glycans, e.g. through use of polysaccharide capsule in glycoconjugate vaccines, has proven highly successful in disease protection, saving millions of lives globally [10]. Also in the context of health-associated microbiota, glycosylation of commensal bacteria contribute to the interaction with the host influencing a variety of homeostatic processes, including epithelial barrier integrity and immune modulation [11]. In contrast to proteins, glycans are not directly encoded by the bacterial genome, but are synthesized by complex biosynthetic pathways involving multiple building blocks and enzymes. This hampers the application of routinely-used omics approaches to capture the microbiota glyco-landscape. Similarly, glycan-microarrays are a useful technique for studying glycan-protein interactions, yet they are unable to capture the diversity of the microbiota glycome at a species-specific level [12,13]. Consequently, the microbiota glycome and its impact on human health and disease represents a major knowledge gap.

Coupling fluorescence-based cell sorting with genomic sequencing has proven a powerful pipeline for identifying subsets of microbiota species with specific immunological properties. For example, IgA-seq allowed identification and subsequent characterization of colitogenic bacteria in patients suffering inflammatory bowel disease from based on differential IgA coating [14,15]. Similarly, two recent studies used glycan-binding lectins as probes to identify microbiota species or even strains with specific glycan patterns. The Glycan-seq technology used 39 different DNA-barcoded non-mammalian lectins to compare glycan profiles of adult mice and pup microbiota samples [16]. The Lectin-seq technology showed the selective and distinct labeling of commensals by two soluble human innate lectins, mannose-binding lectin and intelectin-1, to human fecal microbiota species [17]. In addition to soluble lectins, the repertoire of innate immune receptors also encompasses a wide range of cell-expressed C-type lectin receptors (CLRs), which are abundantly expressed by antigen-presenting cells that line the mucosal surfaces and sample the environment through trans-barrier sampling [18-20].

In this study, we aimed to study the interactions between the bacterial gut microbiota and APC-expressed CLRs. To this end, we have developed a pipeline, called CLR-sequencing (CLR-seq), which combines fluorescence-based sorting of CLR-reactive bacteria with 16S rRNA gene sequencing. As a proof-of-concept, we used two human CLRs with distinct glycan specificities, i.e. the macrophage galactose-like C-type lectin (MGL; CD301), which predominantly interacts with terminal *N*-acetylgalactosamine (GalNAc) residues, and langerin (CD207), which binds high mannose structures, fucose, *β*-glucan and *N*-acetylglucosamine (GlcNAc)[21,22]. Both CLRs recognize distinct glycan moieties on the cell wall of specific bacterial pathogens such as *Staphylococcus aureus*. Moreover, the CLR-bacteria interaction modulates the maturation and cytokine expression of the interacting APC [23,24]. Using langerin and MGL, we show the ability of these CLRs to identify *S. aureus* in bacterial mixtures based on their expressed glycan profile. Next, we applied the two CLRs to fecal microbiota samples from three healthy human donors to identify microbiota species that express MGL and/or langerin-reactive glycans. We envision this approach to open new avenues to study the interaction between host immune system and microbiota glycome and uncover interaction that are relevant to human health and disease.

## Results

### Soluble CLRs identify bacteria with specific glycan motifs in synthetic mixtures

The aim of this study was to establish a pipeline for identification of bacterial microbiota species that can be recognized by human APC-expressed CLRs. For this proof-of-concept study, we used the macrophage CLR MGL (CD301) and the Langerhans cell CLR langerin (CD207), which have distinct glycan binding specificities [21,22]. First, we validated that our approach could selectively identify and extract bacteria based on their surface glycan composition from a mixed bacterial community. To this end, we created a synthetic microbial community of equal proportions of four bacterial species, i.e. two Gram-positive (*Streptococcus thermophilus* and *Listeria monocytogenes*) and two Gram-negative bacteria (*Escherichia coli* and *Alcaligenes faecalis*) that did not bind to either recombinant human langerin or MGL (Figure 1a). We then added CLR-binding *S. aureus* bacteria to this mixture in a proportion of 20% relative to the total community. For MGL, we used *S. aureus* PS187, which expresses MGL-binding GalNAc attached to surface-anchored wall teichoic acid (WTA) [23]. *S. aureus* strain N315 was used to validate langerin-based identification and sorting as this strain specifically binds through the β-linked GlcNAc on WTA [24].

**Figure 1.**
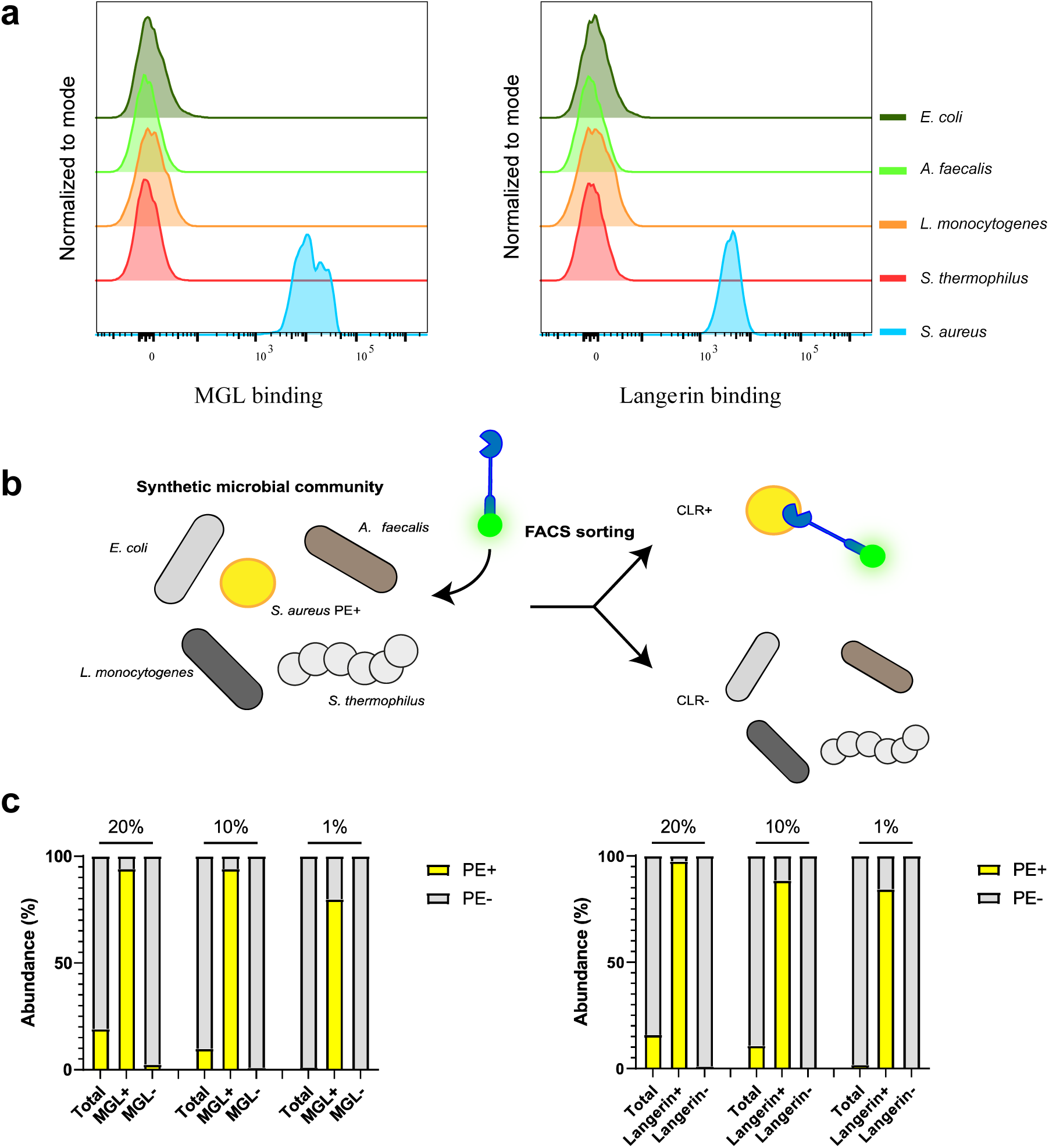
CLRs can identify bacteria with specific glycan patterns in a synthetic microbial community. **a** recombinant human MGL or langerin-FITC staining of the four bacterial species that were used to create a synthetic microbial mixture. *S. aureus* PS187 and N315 are included as CLR-binding positive controls. **b** Cell trace yellow (CTY)-stained *S. aureus* N315 or PS187 (identified in PE channel) were spiked into the synthetic microbial community and the mixed population was subsequently stained with either recombinant human MGL or langerin-FITC. The CLR-positive-and negative populations were sorted. **c** Proportion of CTY-stained *S. aureus* in the total, CLR-positive and CLR-negative sorted populations.

Both *S. aureus* strains were labeled with Cell Trace Yellow (CTY) before mixing the strains, allowing identification of these bacteria in the phycoerythrin (PE) channel and discriminate it from the synthetic bacterial mixture (Figure S1). Subsequently, the bacterial mixture was stained with FITC-labeled his-MGL or langerin and FITC-positive and -negative bacteria were sorted using fluorescence-based cell sorting (Figure 1b). Re-analysis of sorted CLR-positive and -negative populations showed strong enrichment of CTY-stained *S. aureus* in the CLR-positive population and depletion in the CLR-negative populations (Figure 1c). These results show that fluorescence-based cell sorting of bacteria based on CLR-staining is glycan specific.

Bacterial species are present at different abundances in gut microbiota. Therefore, we aimed to show that CLR-based cell sorting would allow identification of low-abundant bacteria in a complex synthetic microbial community by decreasing the proportion of CTY-stained *S. aureus* PS187 or N315 cells to 10% and 1% of the total synthetic bacterial community. Similar to previous results, bacteria could be discriminated from the synthetic bacterial community by double staining even at lower abundance (Figure S1). Sorting and re-analysis of CLR-positive and -negative bacteria showed that independent of the proportion of *S. aureus* added to the mixture, the negative population was completely depleted of CTY-positive *S. aureus* (Figure 1c). Complementary, even at 1% abundance, the positively-sorted fractions were strongly enriched for *S. aureus,* with approximately 80% CTY-positive cells at 1% presence (Figure 1c). Taken together, these data indicate that bacterial strains with specific glycan motifs can be identified and sorted from more complex microbial communities.

### CLR-sequencing on human microbiota

We have thus far only stained and identified known CLR-binding *S. aureus* strains in a synthetic community of limited diversity. To identify potentially new bacterial species that are recognized by human MGL or langerin, we coupled CLR-based binding and sorting to 16S rRNA sequencing. We first stained isolated human fecal microbiota with the DNA stain SYTO60 to determine the bacterial presence and purity of our isolation protocol. On average, 85% of events was SYTO60 positive (Figure S2), implying that our isolation protocol removes most debris and non-bacterial particles. Upon incubation of the isolated fecal material with FITC-labeled MGL or langerin, we observed that both CLRs bound to a subset of the human microbiota (Figure 2). Addition of the calcium chelator EGTA abrogated binding (Figure 2), which confirms the calcium-dependent nature of the CLR interactions [25]. Next, the CLR-binding FITC+ populations were sorted and collected. Re-analysis of the CLR-positive sorted fractions showed 89% and 94% SYTO+ events for Langerin and MGL, respectively, indicating the high presence of bacterial DNA (Figure S2).

**Figure 2.**
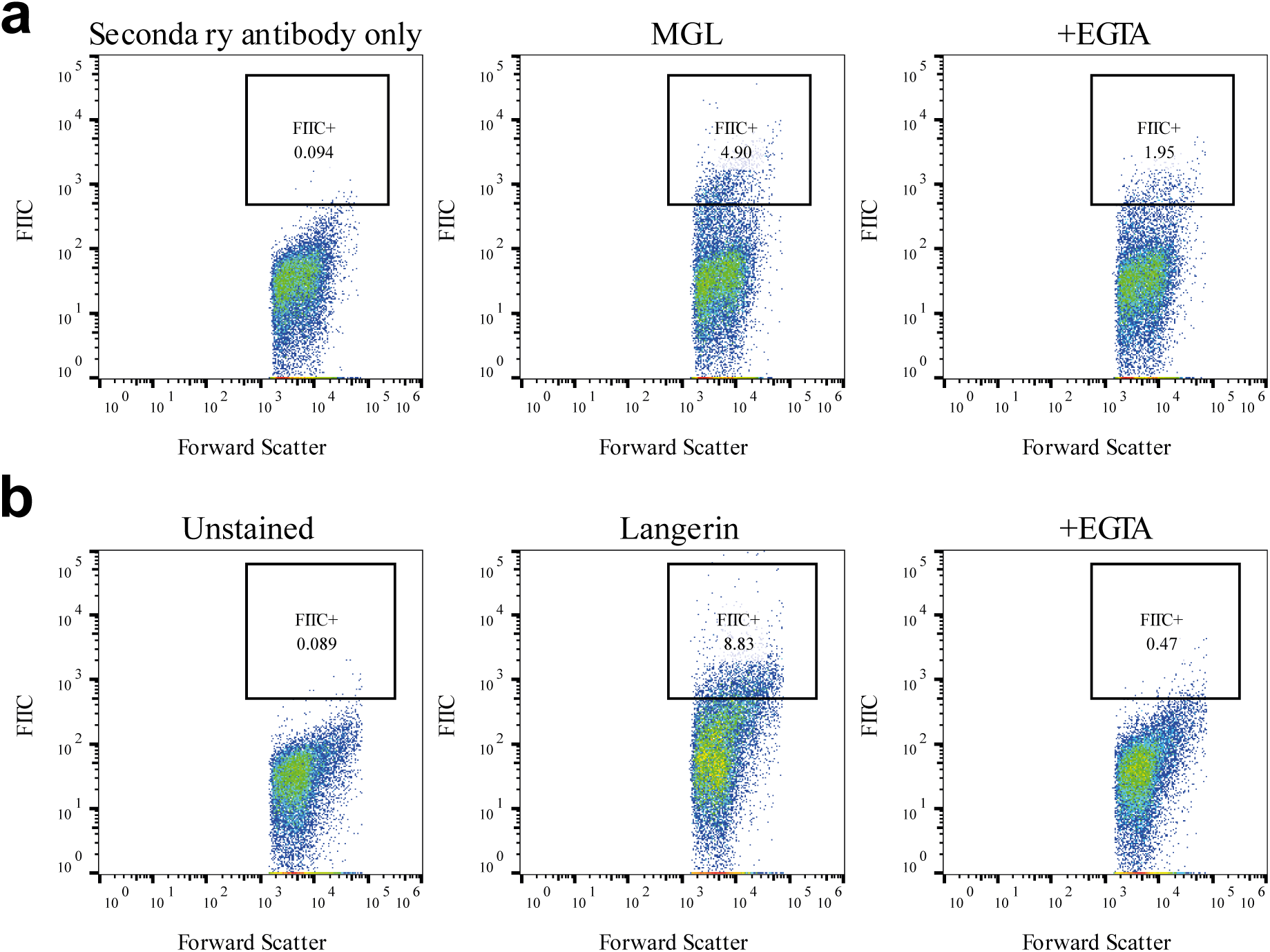
Human CLRs bind to specific microbiota populations in a calcium-dependent manner. Isolated human fecal microbiota was stained with recombinant human **a** MGL or **b** langerin (middle panels), resulting in FITC-positive populations as compared to either secondary antibody only or unstained samples (left panels). CLR binding was abrogated by the addition of the calcium chelator EGTA (right panels).

Having established that our pipeline allows identification of CLR-reactive bacterial strains/species with little non-bacterial impurities, we applied the CLR-seq pipeline to isolated fecal microbiota samples from three individual healthy human donors. Fecal microbiota samples were incubated with MGL and langerin and positive fractions were sorted using fluorescence-activated cell sorting. Additionally, we identified and sorted IgA-positive bacteria from the same fecal samples as it was previously shown that this pipeline allows identification of bacterial species with immunostimulatory properties. We observed inter-individual variation in the proportion of IgA-, MGL- and langerin-reactive bacteria of the healthy donors, where the proportion of CLR-reactive bacteria was smaller compared to the IgA-positive fraction (Figure 3a). Additionally, we observed inter-individual variation in these proportion of MGL-, Langerin- and IgA-reactive bacteria of the healthy donors.

**Figure 3.**
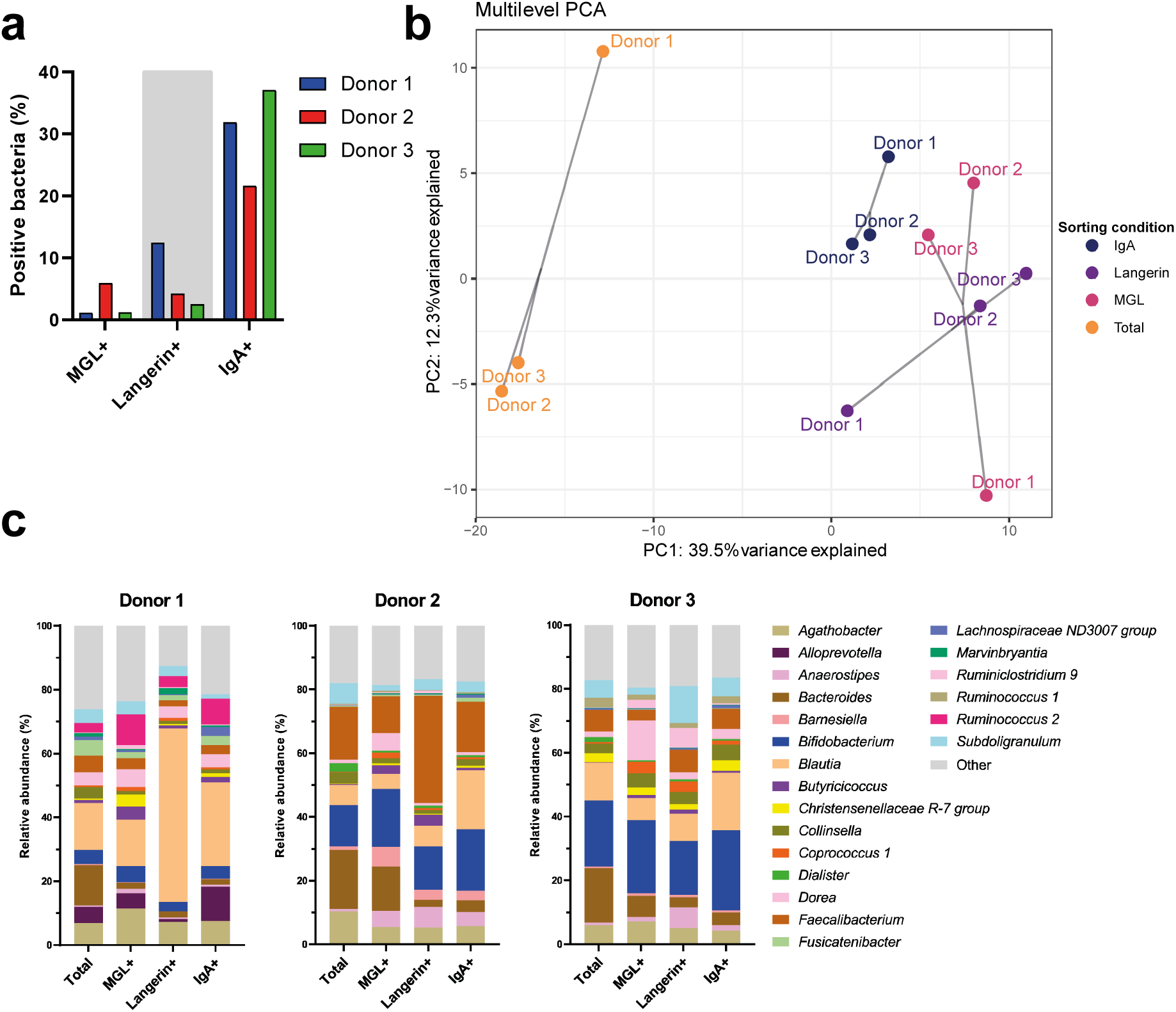
Characteristics of CLR- and sIgA-positive populations from fecal microbiota. **a** MGL-, langerin and sIgA-positive bacterial populations were collected from fecal microbiota samples (n=3 healthy donors), using fluorescence based cell sorting, and analyzed with 16s rRNA sequencing. **b** Principle component analysis of all sequenced samples after multilevel decomposition for repeated measurements in individual donors**. c** Relative abundances at the genus level for genera that were among the 10 most abundant genera in at least one sequenced sample.

Subsequently, DNA was extracted from the total input as well as IgA- and CLR-positive sorted fractions and analyzed by 16S rRNA gene sequencing to identify the bacteria species in the enriched fractions. We observed a decrease in species diversity based on the Shannon Diversity index post-sorting when compared to the unsorted, total populations (Figure S3a). Importantly, the sequencing data varied most strongly between the total unsorted microbiota samples and the sorted IgA and CLRs fractions based on a principal component analysis (PCA) with multilevel decomposition for repeated measurements (Figure 3b). Based on the 16S rRNA gene sequencing, relative abundances were determined at various taxonomic levels for the total populations and sorted samples. Despite interindividual variation between donors at the genus taxonomic level, some shared changes in relative abundances could be observed (Figure 3c). For all three donors, the relative abundance of *Bacteroides* bacteria was decreased for MGL-, langerin-, and IgA sorted samples. In contrast, the proportion of *Butyricicoccus* and *Coprococcus 1* was increased in all three MGL sorted donor samples. The langerin samples for donor 1 and 2 showed an increase in relative abundance for *Blautia* and *Faecalibacterium* bacteria, respectively. When examining the sample composition at the taxonomic species level, we observed a large presence of specifically *Blautia obeum* in donor 1 and an *Faecalibacterium sp.* (zOTU_579) in donor 2 of the langerin sorted samples (Figure S3b).

Using the relative abundances from the 16S rRNA gene sequencing analysis, positive probability scores were calculated for all zOTUs as described previously [26]. To avoid the introduction of potential contaminants during sample preparation, collection or sequencing, we only included zOTUs that were detected in the total population of all three donors and had a relative abundance of >0.1% in at least two donors. Probability scores of zOTUs in the MGL-, langerin-, or IgA positive samples did not correlate with their relative abundance in the total population for any of the donors, suggesting that species abundance in the unsorted microbiota population does not influence CLR- or IgA-binding (Figure S4).

Plotting the individual values for the zOTUs with the highest and lowest average probability scores showed the individual differences in binding of specific species (Figure 4). In the zOTUs with the highest average MGL binding (Figure 4a), donor 2 often showed the highest score, whereas for langerin (Figure 4b) and IgA (Figure S5) this varied more among the donors. For both MGL and langerin, the zOTUs with the lowest average positive probability scores consisted mostly of *Bacteroides sp.* with a number of these being completely depleted in the sorted sample, as indicated by a positive probability score of 0.

**Figure 4.**
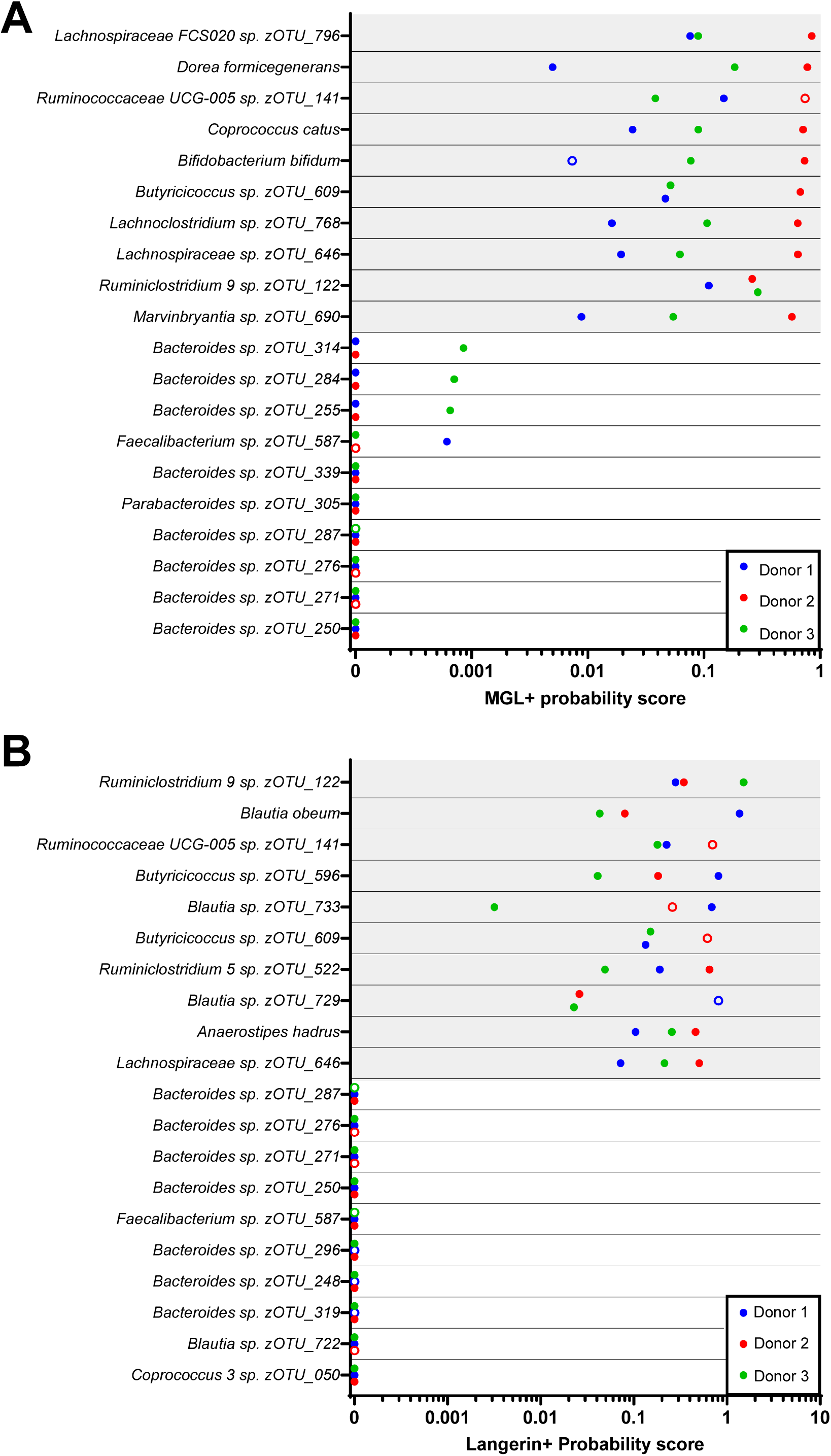
CLRs bind to specific human gut bacteria species. Probability scores were calculated for CLR binding based on the 16s rRNA sequencing relative abundances. Shown are the 10 zOTUs with either the highest (grey background) or lowest average probability scores (white background) for individual donors for **a** MGL or **b** langerin. Open circles are used for donors where the zOTU had <0.1% relative abundance in the total population.

*Marvinbryantiasp sp. zOTU_690* and *Ruminococcaceae UCG-005 sp. zOTU_141* were among the zOTUs with the highest average probability scores for MGL and langerin. Interestingly, these species also showed high sIgA binding (Figure S5). Complementary, zOTUs with low probability scores for CLR binding also had low sIgA probability scores (Figure S5). Including all zOTUs in the sorted fractions, the average probability scores of the CLR- and IgA-positive zOTUs correlated significantly (Figure S6).

### Validation of CLR binding to species identified by CLR-seq

Interestingly, we observed *Blautia sp.* zOTUs in both the highest and lowest average langerin probability scores. This indicates that CLR-seq can discriminate within the taxonomic genus level between species expressing different glycan profiles [27]. We therefore deemed it important to validate a number of species in the CLR-binding fraction as monoculture (Figure S7).

*Bifidobacterium bifidum*, a gut commensal with probiotic characteristics and present in the intestinal tracts of nearly every individual [28], and *Dorea formicegenerans*, another intestinal commensal bacterium, were among the species with the highest average probability scores for MGL (Fig. 4A). To validate the CLR-seq results of these donors, MGL binding was tested on monocultures of a *B. bifidum* and *D. formicegenerans* strain. MGL interacted with *B. bifidum* and this binding was dependent on the carbohydrate recognition domain (CRD) of MGL and bivalent cations, since MGL binding was abrogated by the addition of GalNAc and significantly reduced in the presence of EGTA, respectively (Figure 5a). No binding of MGL to the *D. formicenegerans* monoculture was observed.

**Figure 5.**
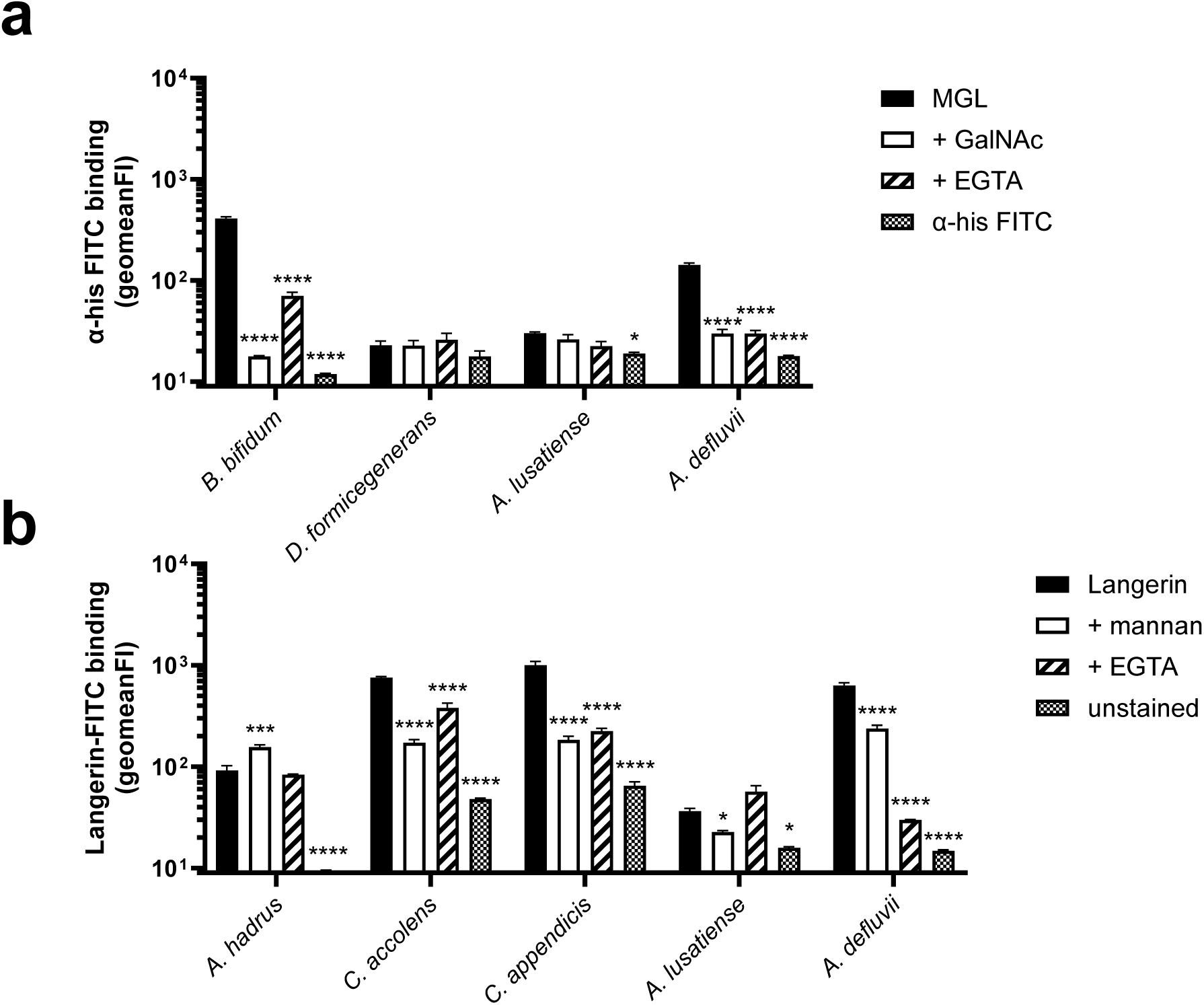
Validation of CLR binding to bacterial species identified through CLR-seq. Binding of **a** recombinant human MGL and **b** recombinant human langerin-FITC to bacterial species identified by CLR-seq. CLRs were used at a concentration of 25 µg/mL, similar to the concentration used during sorting. Binding was blocked using *N*-acetyl-galactosamine (GalNAc) or mannan for MGL and langerin, respectively, and with the calcium chelator EGTA. Statistical comparisons indicated are the blocked or unstained bacteria compared to the stained control. Data are shown as the mean of the geometric mean fluorescence intensity (FI) ± standard error of mean from three independent experiments. *, P<0.05; ***, P<0.001; ****,P < .0001

For langerin binding, the gut commensal bacterium *Anaerostipes hadrus* was among zOTUs with the highest average probability scores (Figure 4b). Therefore, we tested binding of this CLRs to a *A. hadrus* monoculture (Figure 5b). We observed langerin-FITC binding, although this binding was not abrogated by the addition of EGTA, and even increased by the addition of mannan. These data suggest that this binding to our recombinant langerin construct was a-specific.

To avoid the interference of contaminants in our 16s rRNA sequencing analysis, we had removed zOTUs that were not detected in the total population of their respective donor. However, when examining the removed zOTUs more closely, we observed consistent presence of various zOTUs in the CLR-positive fractions of our three donors. For both MGL and langerin-positive fractions, we detected the genus *Aquamicrobium*, which was first described in activated sewage sludge [29]. In our 16S rRNA gene sequencing data, this genus consisted of a single zOTUs attributed to two species, *A. lusatiense* and *A. defluvii*. Additionally, we identified two zOTUs at the species level within the genus *Corynebacterium* with consistent presence in the langerin-positive fraction. These were *C. accolens*, an abundant nasal commensal, and *C. appendicis*, which was first isolated from the abdominal swab of an appendicitis patient [30,31]. To verify CLR binding to the mentioned species, an individual strain of each species was grown as a monoculture and incubated with recombinant human MGL or langerin, corresponding to the sorted fractions in which they were identified (Figure 4). MGL bound significantly to both *A. defluvii* and *A. lusatiense*, although this binding was higher in *A. defluvii*. Furthermore, for *A. defluvii*, but not *A. lusatiense*, this binding was abrogated by the addition of GalNAc and EGTA (Figure 5a), suggesting that the recognition by MGL occurred through the CRD and was calcium dependent in this bacterium. Similarly, we observed significant langerin binding to both *Aquamicrobium* species, with *A. defluvii* showing higher binding and a more significant loss of binding by the addition of mannan or EGTA (Figure 5b).

*C. appendicis* and *C. accolens* both bound to recombinant human langerin in a CRD and calcium dependent manner (Figure 5b), although the binding appeared to be saturated at a lower concentration for *C. appendicis* (Figure S7b). These results highlight the potential of CLR-seq to specifically identify bacteria that can be recognized by CLRs.

## Discussion

Bacterial glycans are critical for both cell viability, as well as interaction with the host-immune system. However, the complex nature of the biochemical pathways involved in glycan synthesis complicates glycan research on the level of complex microbial communities, such as the gut microbiota. Previously, microbiota glyco-profiling was performed using a large panel of non-mammalian lectins and two soluble human lectins [16,17]. Here we analyzed the human microbiota through C-type lectin receptor-sequencing (CLR-seq), which combines interaction with human surface-expressed CLRs with 16S rRNA sequencing. We present proof-of-principle that CLR-seq can be used as a platform to comprehensively determine the microbiota-binding capabilities of human CLRs. The approach can easily be adapted to other host CLRs or microbiota samples of interest, for example from different mucosal sites or from patients with a specific clinical condition.

To establish proof-of concept for CLR-seq, we used MGL, which is expressed by tissue resident macrophages and langerin, a Langerhans cell-specific CLR, since we have extensive experience with these two CLRs in host-microbe interaction [23,24]. We chose to apply CLR-seq on fecal microbiota samples since this is the most abundant microbial community present in the human body, thereby yielding the most diverse glycome. Currently, we have limited insight into the exact APC subsets and associated CLRs that line the different intestinal barriers in humans. In mice, the presence of the murine MGL homologue was found in colon tissue [32]. Additionally, MGL expression was found on APCs located in the lamina propria of the jejunum [33]. Although langerin is most likely not present in the gut, the CLR DC-SIGN, predominantly expressed by dendritic cells, was identified in human Peyer’s patches [34] and shows substantial overlap in ligand binding with langerin [refs]. The incorporation of additional CLRs, such as DC-SIGN, would give a more complete overview of the glycan binding capabilities of the mucosal immune system of the gut.

We showed that CLR-seq can specifically target bacteria based on their glycan moieties, even at low abundance in a synthetic microbial community. We then applied CLR-seq to the fecal microbiota of three healthy human donors. To identify bacteria in the unsorted and CLR-sorted fractions, we used 16S rRNA gene sequencing. This posed limitations for the downstream workflow. While we obtained general taxonomic information with a limited bacterial input (200,000 positive events sorted), identification at the species level was not possible for most zOTUs, which was also observed to a lesser extent at the genus level. Since bacterial surface composition of glycans can be strain specific or even phase variable [6], this could result in the unsuccessful validation of certain species. Indeed, we observed that within a genus, individual zOTUs could have either high or low CLR-probability scores. Furthermore, as CLR-seq studies the host-microbe interaction at the native level of a complex bacterial community, this could further contribute to disparities between the identified species in the sorted sequencing data and the monocultures in the laboratory setting. Indeed, these factors highlight the necessity for laboratory validation of identified species. An approach with increased resolution, such as next generation whole metagenome sequencing, could provide a more detailed insight into the bacterial identity and characteristics that confer interaction with innate CLRs. Strain-level information may also resolve discrepancies for species that are inconsistently identified in microbiota screens or are being associated with certain host pathologies.

For the zOTUs we identified in our CLR-binding fractions, we could validate MGL binding to *Bifidobacterium bifidum*, a human commensal, in a CRD-dependent manner. MGL predominantly binds to galactose and GalNAc residues, which are present in certain *B. bifidum* polysaccharide structures [35]. The non-specific langerin binding to *A. hadrus* and lack of MGL binding to *D. formicegenerans* may be related to the factors described above, i.e. strain variation and environmental conditions that affect glycan patterns and expression.

We observed a significant correlation between CLR- and IgA binding for the microbiota species identified by 16s rRNA sequencing. Interestingly, it has been described that affinity-matured microbiota-binding sIgA is often glycan-reactive [36,37]. We therefore speculate that CLRs and sIgA may target similar glycan structures. Alternatively, there is the possibility that the used CLRs bind to glycan moieties attached to sIgA [38], which selectively binds to a specific subset of gut microbiota species. However, the CRD-dependent binding of *B. bifidum* as a monoculture shows that this is certainly not the case for all CLR bound bacteria. Additionally, we could validate binding to species that were suspected environmental contaminants. These species were not detected in the unsorted microbiota populations, but were consistently present in the CLR positive populations. This suggest the selectivity of CLR-seq even at very low abundances and independent of sIgA ccoating.

In the context of infection, CLRs expressed on APCs have an important role of sensing a wide array of pathogens, including viruses, bacteria and fungi, through interaction with specific microbial glycan patterns that are abundantly expressed at the surface [18,39,40]. But also during homeostasis, APCs constantly probe the local microbiota and bacterial antigens at host-barrier sites by extending their dendrites into the lumen [5,19,20]. This sampling and further processing contributes to the development of local and systemic immunity that help maintain homeostasis but also clearance of invading pathogens [41,42]. Interstingly, CLRs are densely expressed on the probing dendrite tips [43,44]. Hence, CLRs on APC dendrites are uniquely equipped and localized to selectively sample bacteria from a complex microbiota community based on their glycan specificity during homeostasis. As an outlook, we therefore anticipate that CLR-seq may provide a greater understanding of microbiota species selection by glycan-dependent innate immune sampling by mucosal APCs.

Overall, our results demonstrate that CLR-seq can identify bacterial microbiota species based on their glycan pattern or sIgA binding from complex microbial communities. This technique can easily be expanded to an increased panel of recombinant immune receptors, the incorporation of samples from different mucosal compartments, as well as the comparison of the microbial glycan content of healthy individuals with patients with underlying disease, making it a new tool for functional microbiota research.

## Methods

### Fecal donor samples

The fecal donor samples used were collected as part of the PIMMS study (NL67136.018.18) performed at Amsterdam UMC (Amsterdam, the Netherlands) as previously described [45]. Fecal matter was collected from healthy participants and stored at -80°C until further processing. All included donors provided informed consent and the study was approved by the Ethical Review Board of the Amsterdam UMC.

### Bacterial strains and culture conditions

The bacterial strains used in this study are listed in Table S1. *Staphylococcus aureus* and *Aquamicrobium* strains were grown in Tryptic Soy broth (TSB; Oxoid) and *Escherichia coli* in Luria Bertani (LB; Oxoid) broth. *Streptococcus thermophilus* was grown in Todd-Hewitt broth supplemented with 1% yeast extract (THY, Oxoid) in the presence of 5% CO_2_ and *Listeria monocytogenes* in Brain Heart Infusion (BHI, Oxoid) broth. Overnight cultures were diluted to an optical density 600 nm (OD600) of 0.05 in fresh medium and grown to mid-exponential phase, after which bacteria were harvested by centrifugation. *Alcaligenes faecalis* and the *Corynebacterium* species were grown on Columbia blood agar (Oxoid), with *Corynebacterium appendicis* grown in a microaerophilic environment. Bacteria were collected from plate using an inoculation loop, and washed with Tris-sodium-magnesium buffer (TSM: 20 mM Tris; Roche, 150 mM NaCl; Merck, 2 mM CaCl_2_; Merck, 2 mM MgCl_2_; Merck, pH 7.0) + 0.1% Bovine Serum Albumin (BSA). *Bifidobacterium bifidum* and *Dorea formicegenerans* were isolated from human feces as described previously [14]. Subsequently, they were grown in an anaerobic chamber in Gut Microbiota medium (GMM) [46]. Overnight cultures were diluted in fresh medium to an OD600 of 0.05 and grown for 6 hours, after which they were harvested by centrifugation. *Anaerostipes hadrus* was cultured in modified yeast extract, casitone and fatty acid (YCFA) medium [47] supplemented with 30 mM glucose. Overnight cultures were diluted in fresh medium (1:10) and grown for 4 hours, after which they were harvested by centrifugation. All species were grown at 37°C, except for the *Aquamicrobium* species, which were grown at 30°C.

### Expression and purification of recombinant human MGL

The extracellular domain of human MGL (CLEC10A) was expressed and purified as described previously [48]. A gBlock encoding the open reading frame of human MGL and a C-terminal LPETGG-6xHis tag were cloned into a pcDNA34 expression vector (Invitrogen). Expression of recombinant hMGL-his proteins was performed in Expi293F cells (Life Technologies), which were cultured in Expi293 Expression Medium (Life Technologies). The protein was purified from the supernatant using affinity chromatography (ÄKTA Pure, GE Healthcare Life Sciences) using a Nickel column (GE Healthcare Life Sciences) as previously described (Zhao et al, 2020). Eluate was dialyzed against 300 mM NaCl 50 mM Tris pH 7.8 at 4°C.

### Staining and flow cytometric analysis of bacterial strains

Bacteria were collected from broth culture by centrifugation or from plate with an inoculum loop after overnight growth, washed with and resuspended in TSM + 0.1% BSA after centrifugation (3,600 x g, 10 min, 4°C). Bacterial abundance was determined by optical density at 600 nm or using flow cytometry with Precision Count Beads (Biolegend 424902) using a BD FACSymphony or BD FACSCanto.

For langerin and MGL specificity analysis in synthetic microbial communities, *S. aureus* strains N315 and PS187 (Table S1) were used as positive controls, respectively [23,49]. Both *S. aureus* strains were fluorescently-labeled with Cell Trace Yellow (CTY; Invitrogen) as per manufacturer’s instructions and mixed at desired relative abundances in the synthetic microbial community consisting of 10^7^ bacterial cells (quantified by flow cytometry). Subsequently, the bacterial mixture was stained with either FITC-labeled recombinant langerin (kindly provided by Prof. Christoph Rademacher, University of Vienna, Austria) or his-tagged recombinant human soluble MGL at 25 µg/mL in TSM + 0.1% BSA for 30 min at 37°C (900 rpm shaking, covered from light). After washing with TSM + 0.1% BSA, hMGL binding was detected using an anti-hisTag FITC-conjugated antibody (1:20 in TSM + 0.1% BSA, Invitrogen; clone AD1.1.10), incubated for 20 minutes at 4°C, covered from light. FITC-positive and - negative bacteria were sorted and analyzed using a Sony SH800 flow cytometric cell sorter.

To determine binding of CLR constructs to bacterial monocultures, 10^6^ CFU (determined by OD600) or bacterial cells (quantified by flow cytometry) were stained with langerin-FITC or his-hMGL as described above. MGL binding was blocked by adding *N*-acetyl-galactosamine (GalNAc; 100 mM; Sigma-Aldrich A2795) and langerin binding was blocked through the addition of mannan (20 µg/mL; Sigma-Aldrich M7504). To test for calcium dependent binding, the calcium chelator EGTA (Sigma Aldrich, 10 mM) was added. After washing with TSM + 0.1% BSA, bacteria were fixed with phosphate buffered saline (PBS) + 1% formaldehyde (Sigma-Aldrich) and measured by flow cytometry (BD FACS Canto).

### Fecal bacteria isolation and staining

Fecal microbiota was isolated as previously described [14] with some modifications. Approximately 100 mg frozen fecal material was resuspended in 1 mL of ice-cold PBS for 15 minutes on ice. Next, the suspension was transferred to a Fast Prep Lysing Matrix D tube containing 1.4 mm zirconium-silicate beads (MP Biomedicals) and homogenized by beat-beating for 10 seconds at 6,000 rpm (MagNa Lyser; Roche). To separate the fecal bacteria from the debris, the homogenized suspension was centrifuged (50 x g, 15 minutes, 4°C) and 100 µL supernatant was transferred to a clean 1.5 mL Eppendorf tube and washed with 1 mL PBS + 1% BSA (8,000 x g, 5 min, 4°C). The fecal pellet was resuspended in 1 mL PBS + 1% BSA and filtered using a 70 µm mini cell strainer (PluriSelect), after which 20 µL was set aside as pre-sort sample.

To determine CLR-specific binding of fecal bacteria, the filtered microbiota suspension was centrifuged and stained with 25 µg/mL recombinant human langerin-FITC or his-tagged human MGL in TSM + 1% BSA, and incubated 30 minutes at 37°C with agitation (600 rpm). MGL-stained bacteria were stained with anti-his-FITC after washing (1:20 in TSM + 1% BSA; Invitrogen, clone AD1.1.10) for 20 minutes at 4°C, covered from light. Samples were washed with and resuspended in TSM + 1% BSA and analyzed by flow cytometric (FACSymphony A1, BD Biosciences) measuring 20,000 events, or sorted based on fluorescent staining using fluorescence-activated cell sorting (Sony SH800). For cell sorting, 200,000 positive events per staining were collected in PBS + 1% BSA, pelleted by centrifugation (8,000 x g, 5 min, 4°C), resuspended in 50 µL PBS and stored at -20°C until DNA extraction.

### DNA extraction, 16S rRNA sequencing

Fecal DNA extraction, 16S rRNA gene sequencing analysis, and bioinformatics analysis were performed as described previously [50]. Briefly, total genomic DNA was isolated from the pre-sorted and sorted microbiota samples at the Microbiota Centre of Amsterdam (MiCA, Amsterdam, the Netherlands). Isolation was performed using a repeated bead beating protocol and the Maxwell RSC Whole Blood DNA kit [51]. DNA concentration was measured in a 96-wells plate using the Qubit dsDNA BR Assay. Amplification of the V3-V4 regions of the 16S rRNA gene sequences was performed with the 341F-805R primers in a single step PCR protocol. Samples were purified using Ampure XP beads, measured and sequenced equimolarly using Illumina MiSeq (v3 chemistry with 2 × 250 cycles).

Amplicon sequences were parsed using a vsearch (v2.15.2) based pipeline [52]. Paired end reads were merged, with max differences set to 100 and allowing for staggered overlap. Zero-radius Operational Taxonomic Units (zOTUs) were inferred from reads with lower than 1.5 expected error rate using the cluster_unoise with centroids algorithm with a minsize of 4, after which chimeras were removed using the uchime3 denovo method. For each sample zOTU abundances were determined by mapping the merged reads against zOTU sequences using usearch_global with a 0.97 distance cut off. Taxonomy was assigned using R (4.2.0) and the dada2 [53] assign taxonomy function using the silva (v132) [54] reference database. A phylogenetic tree was generated use mafft (v7.310) [35] and Fasttree (2.1.11) [55].

The abundance and zOTU tables were used for downstream analysis using R’s phyloseq package [56]. Probability scores were calculated using relative abundances for each genus, similarly as described previously [26] and only genera that were present in the sorted sample of all three individual donors were included. For visualization of 16S rRNA gene sequencing analysis, R-studio (version 4.4.1) or Graphpad Prism (version 10.2.0) was used.

### Statistical analysis

Visualization and statistical analysis were performed using Graphpad Prism (version 10.2.0). Differences in CLR binding at specific concentrations and potential blocking were calculated with a one-way analysis of variance (ANOVA) for each individual species, followed by Bonferroni’s test for multiple comparison. For CLR binding concentration curves, a two-way ANOVA followed by Bonferonni’s multiple-comparison test was used. A P-value of <0.05 was considered significant.

## Acknowledgements

The authors would like to thank Romy Ros and Carla J. C. de Haas for their technical assistance; Dr. Torsten Scheithauer for sharing his expertise with the microbiota sorting process; Jorn Hartman for his help with the preparation of the sequencing library and Professor Christoph Rademacher at the University of Vienna for supplying the recombinant langerin construct.

## Author contributions

JM, RvD and NMvS, planned the experiments. JM performed the experiments and analyzed the data. MEM produced the recombinant MGL construct. MD analyzed the raw sequencing data. MRdZ and HH provided bacterial strains and contributed to study design. JM and NMvS wrote the manuscript with editing by RvD and YP. All authors revised and approved the manuscript.

## Data availability statement

The sequencing data generated in this study have been deposited in the European Nucleotide Archive database under accession code: PRJEB86038. The data are freely available without restriction.

