## Supplemental material for "CLR-Seq: a pipeline to identify bacterial microbiota species with immune-relevant glycan moieties through human C-type lectin receptor interaction"

**Authors:**

Jasper Mol[1], Rob van Dalen[1], Yvonne Pannekoek[1], Malgorzata E. Mnich[2], Marcel R. de Zoete[2], Mark Davids[3], Hilde Herrema[3], Nina M. van Sorge[1,4*]

**Affiliations:**

[1]Department of Medical Microbiology and Infection Prevention, Amsterdam Institute for Immunology and Infectious Diseases, Amsterdam UMC, location University of Amsterdam, Amsterdam, The Netherlands

[2]Department of Medical Microbiology, UMC Utrecht, Utrecht, The Netherlands

[3]Department of Experimental Vascular Medicine, Amsterdam UMC, location University of Amsterdam, Amsterdam, The Netherlands

[4]Netherlands Reference Laboratory for Bacterial Meningitis, Amsterdam UMC, location AMC, Amsterdam, The Netherlands

Corresponding author contact:

Professor Dr. Nina M. van Sorge

Amsterdam UMC, location AMC

Meibergdreef 9

IWO building, room IA3-211

1105 AZ Amsterdam, the Netherlands

**Supplementary Table S1.** Bacterial strains used in this study.

| **Species** | **Strain** | **Source** |
|---|---|---|
| *Staphylococcus aureus* | N315 | Network on Antimicrobial Resistance in *Staphylococcus aureus* (NARSA) strain collection |
| | PS187 | ATCC, Cat#15564, [1] |
| *Escherichia coli* | DC10B | LMBP 9585, [2] |
| *Alcaligenes faecalis* | DSM30033 | German Collection of Microorganisms and Cell Cultures (DSMZ), Braunschweig, Germany |
| *Streptococcus thermophilus* | 2110133 | Netherlands Reference Laboratory for Bacterial Meningitis (NRLBM), Amsterdam, the Netherlands |
| *Listeria monocytogenes* | EGD-E | [3] |
| *Bifidobacterium bifidum* | MSC-333 | Isolated from human feces, this study |
| *Dorea formicegenerans* | MSC-222 | Isolated from human feces, this study |
| *Anaerostipus hadrus* | PEL85 | [4] |
| *Aquamicrobium lusatiense* | DSM11099 | DSMZ |
| *Aquamicrobium defluvii* | DSM11603 | DSMZ |
| *Corynebacterium appendicis* | DSM44531 | DSMZ |
| *Corynebacterium accolens* | DSM44278 | DSMZ |

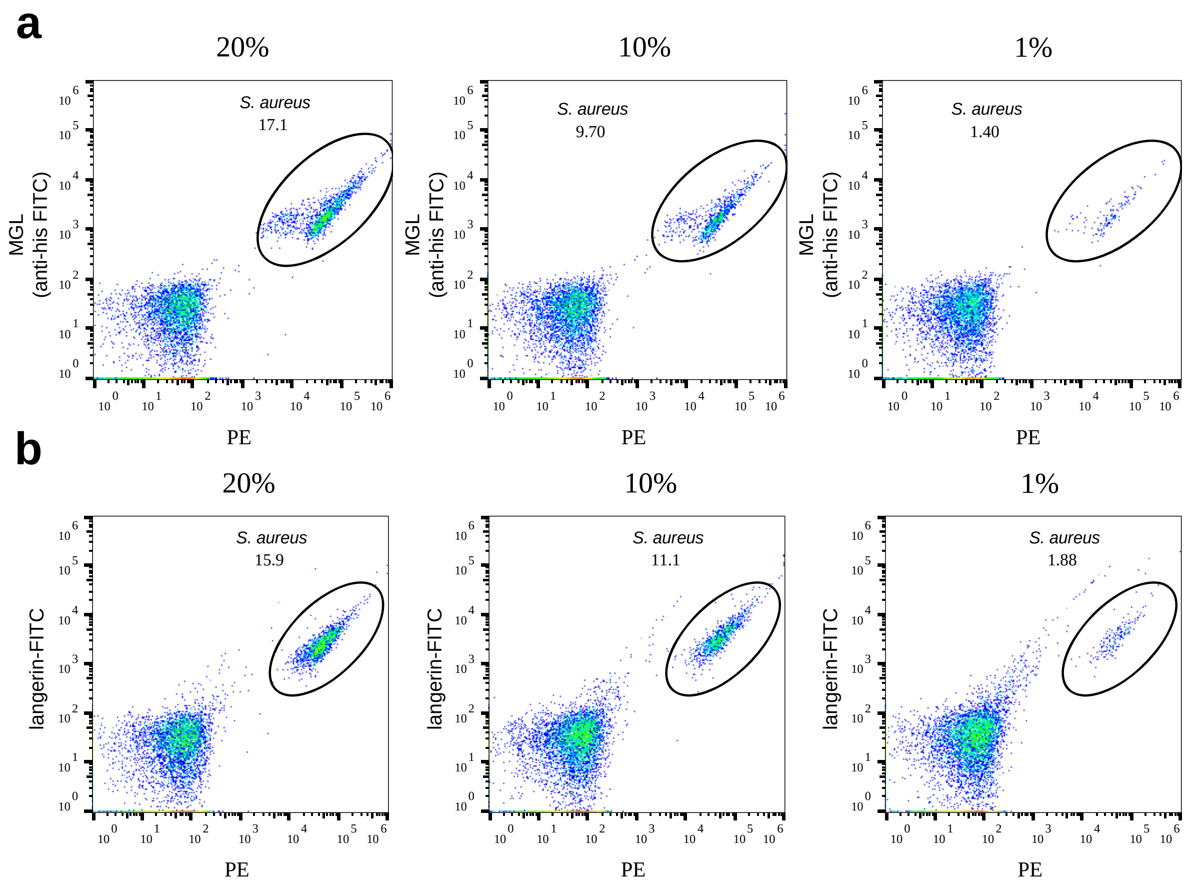


**Figure S1. Soluble MGL and langerin discriminate *S. aureus* in a synthetic microbial community through specific glycan moieties.** Dot blots of a microbial community consisting of equal proportions of *E. coli*, *A. faecalis*, *S. thermophilus*, *L. monocytogenes* and various relative abundances (20, 10, 1%) of CTY-labeled *S. aureus* (visualized in the PE channel). Microbial communities were stained with **a** recombinant human MGL detected with anti-his FITC, labeling only spiked-in *S. aureus* PS187, or **b** langerin-FITC, labeling only *S. aureus* N315. The double-positive *S. aureus* fraction was quantified using flow cytometry.

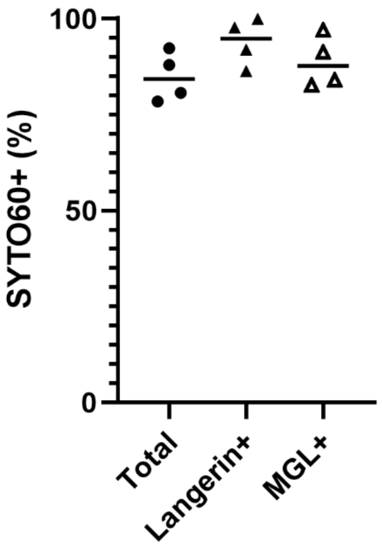


**Figure S2. CLR-sorting pipeline enriches bacterial particles**. Isolated fecal microbiota was stained with the DNA dye SYTO60 to discriminate bacteria from non-animate particles. SYTO60-positive percentages were determined before and after CLR-sorting. Results represent the mean of four individual healthy donor samples.

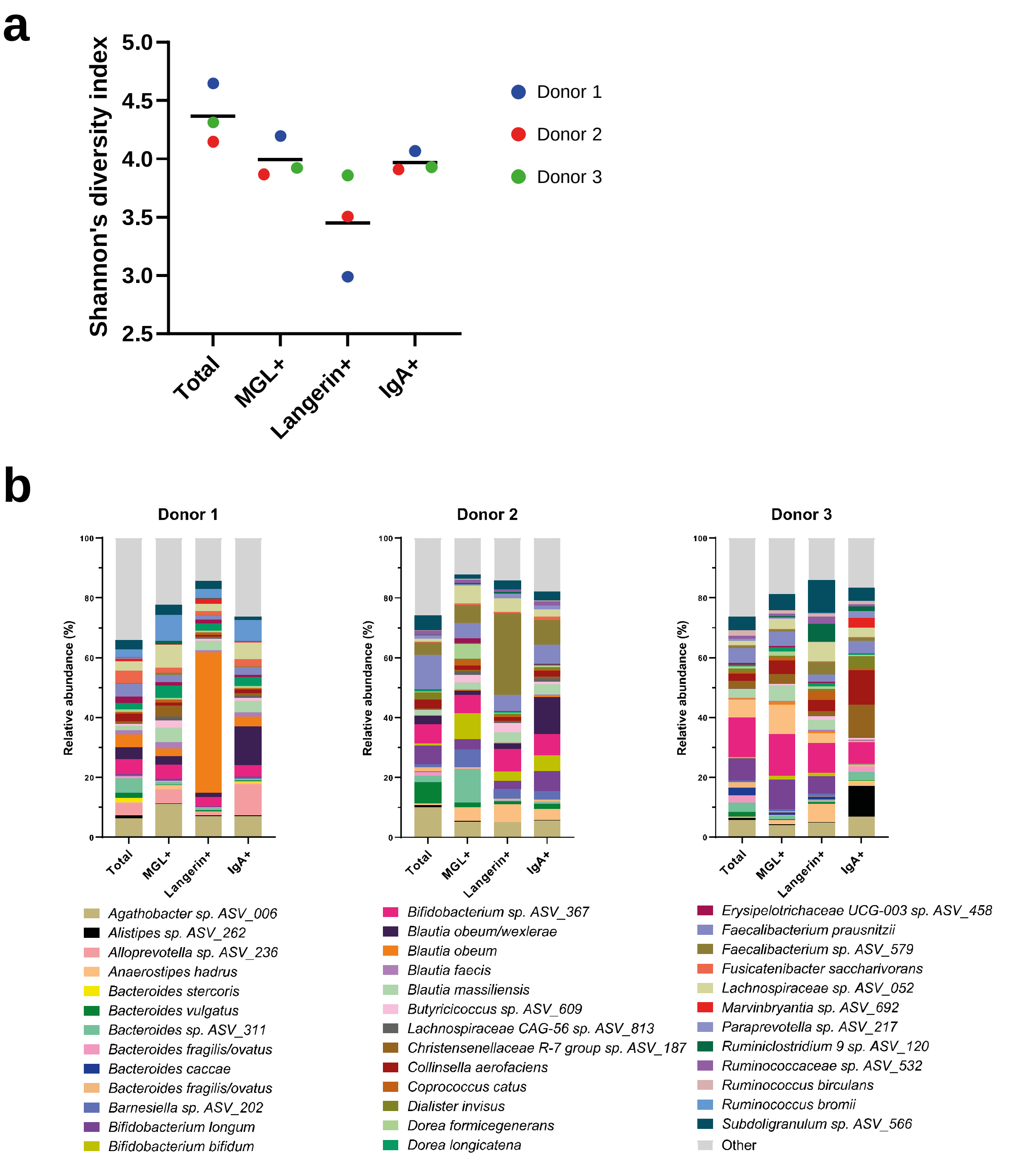


**Figure S3. Changes in bacterial composition after CLR- and IgA-based sorting of fecal microbiota.** **a** CLR- or sIgA-positive bacteria were sorted from fecal samples from three healthy human donors using fluorescence based cell sorting and analyzed with 16S rRNA gene sequencing. The diversity of species was calculated using Shannon's Diversity Index. **b** Relative abundances at the species level of the zOTUs that had a relative abundance of >1% in at least one sorted or unsorted sample across three donors.

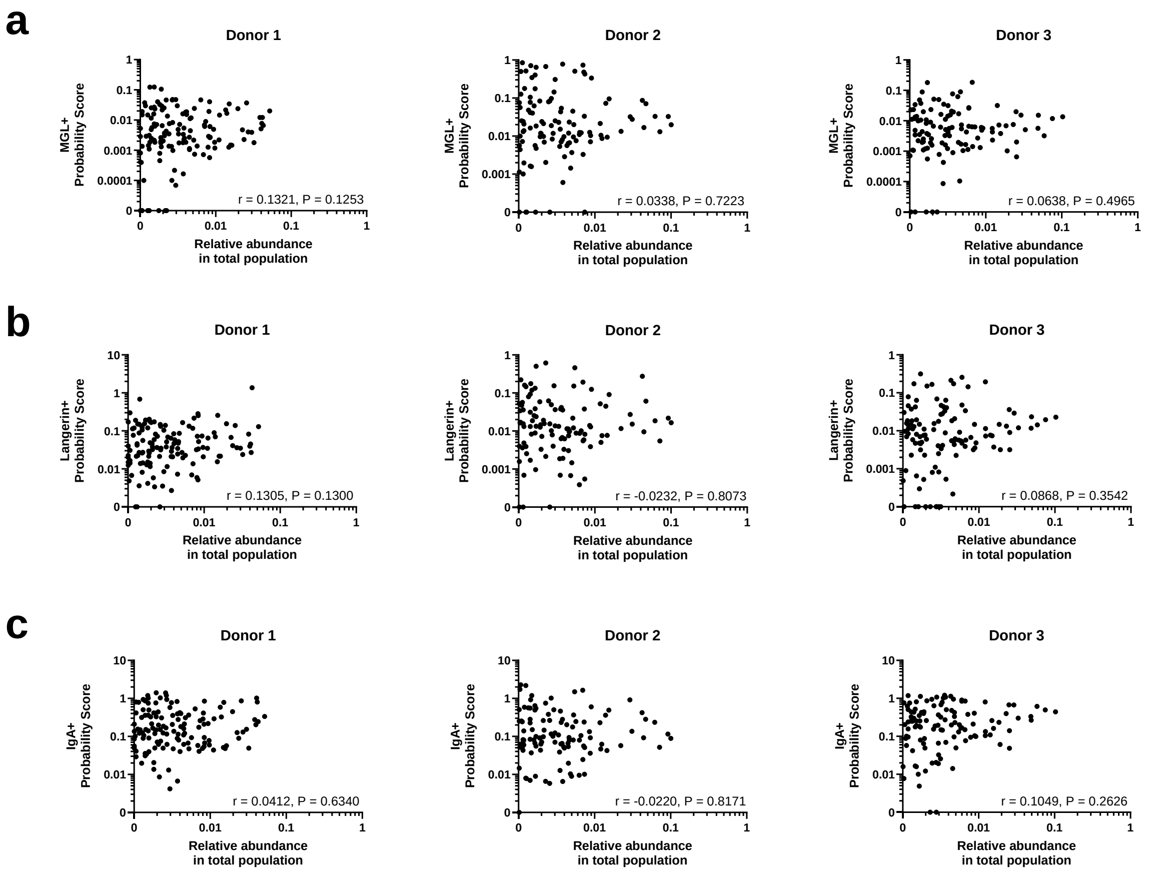


**Figure S4. CLR- and sIgA- binding of microbiota species is not affected by relative abundance in the total population.** CLR- or sIgA binding bacteria were sorted from healthy human fecal donor samples using fluorescence-based cell sorting and analyzed with 16S rRNA gene sequencing. Probability scores were calculated by comparing the relative abundances of identified zOTUs in sorted and pre-sorted (total) populations. Average probability scores for each zOTU in the **a** MGL-sorted, **b** langerin-sorted and **c** **s**IgA-sorted fractions were plotted against their respective relative abundance in the total population. Spearman's rank correlation was used for statistical comparison.

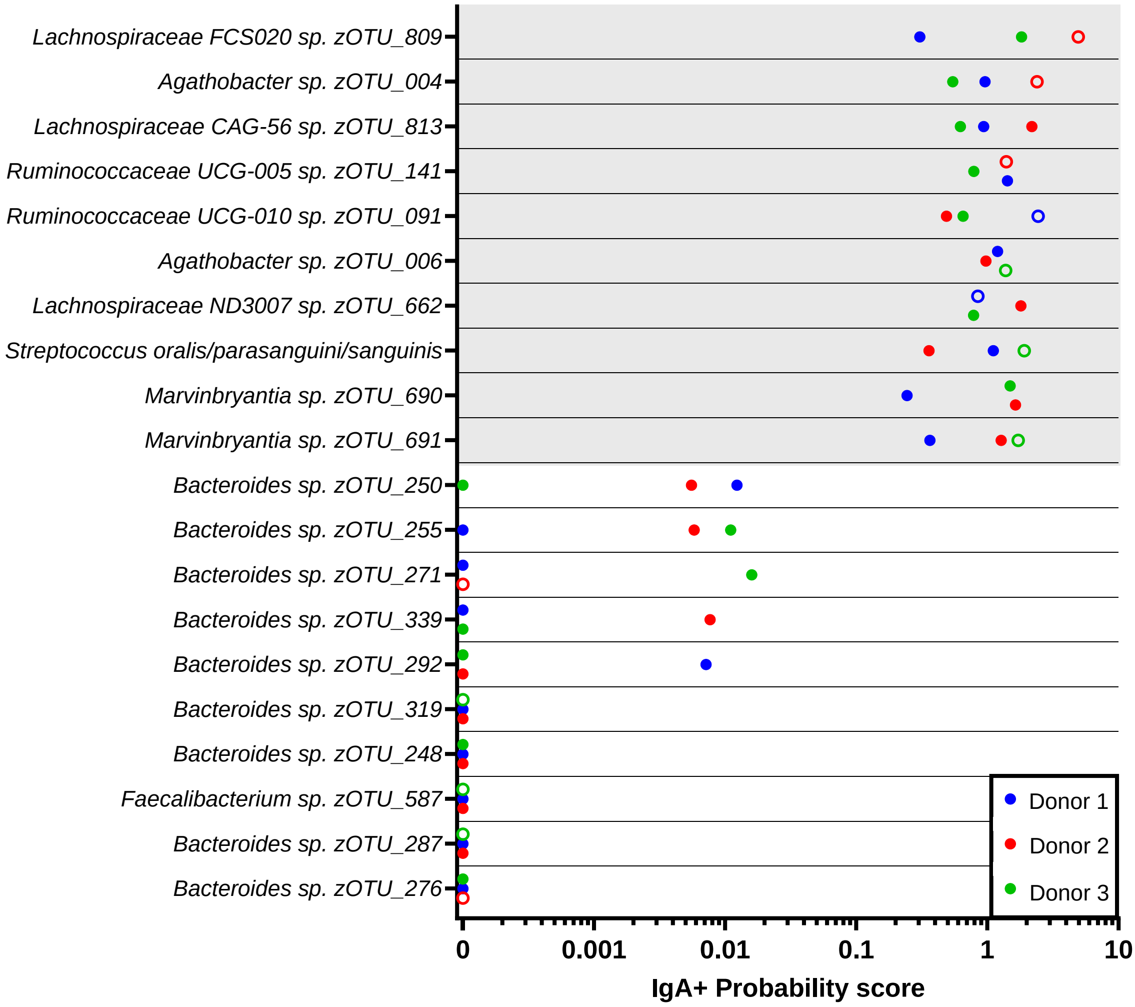


**Figure S5. Identity of sIgA-coated microbiota species based on probability scores.** Positive probability scores were calculated for sIgA binding based on relative abundances in 16s rRNA sequencing data. The graph shows the bacterial species ranking in the top 10 of the highest (grey background) or lowest average probability scores (white background) for IgA binding. Individual scores for each donor are shown. To avoid potential contaminants introduced during sample collection and downstream work up, we only included ASVs that were detected in all three donors and had >0.1% relative abundance in at least two donors in the total (pre-sorted) population. Open circles are used for donors where the ASV had <0.1% relative abundance in the total population.

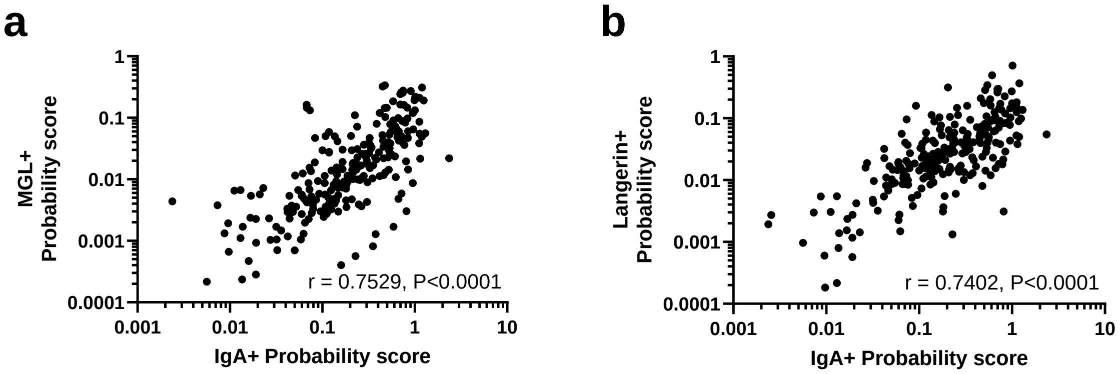


**Figure S6. Correlation between probability scores for CLR-positive and sIgA-coated microbiota species.** CLR- or sIgA-positive bacteria were sorted from fecal samples from three healthy human donors using fluorescence based cell sorting and analyzed with 16S rRNA gene sequencing. Probability scores were calculated by comparing the relative abundances of identified zOTUs in sorted and pre-sorted (total) populations. Average probability scores for each zOTU in the **a** MGL-sorted and **b** langerin-sorted fractions were plotted against their respective sIgA probability scores. Only zOTUs that were present in the total population and had a minimal relative abundance of >0.1% in the total population of at least two individual donors were included (n=251). Spearman's rank correlation was used for statistical comparison.

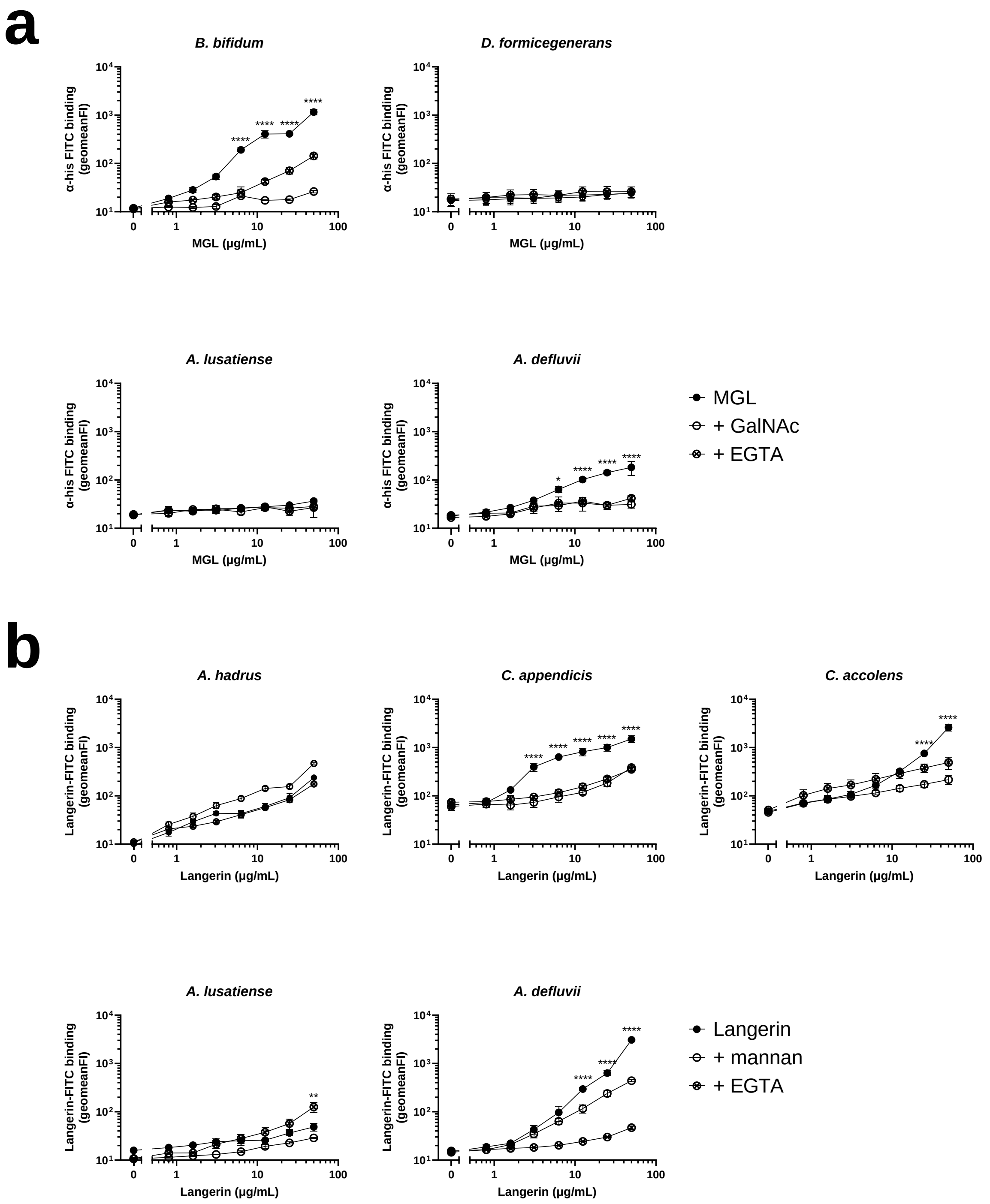


**Figure S7. Validation of bacterial species identified by CLR-seq**. Binding of **a** recombinant human MGL detected by an anti-hisTag-FITC antibody and **b** recombinant human langerin-FITC to bacterial species identified by CLR-seq. CLRs were used at a concentration range (0-50 µg/mL). Binding was blocked using N-acetyl-galactosamine (GalNAc) or mannan for MGL and Langerin respectively and with the calcium chelator EGTA. The indicated statistical differences refer to both blocking conditions compared to the nonblocked control. Data are shown as the mean of the geometric mean fluorescence intensity (FI) ± standard error of mean from three independent experiments. *, $P<0.05$; **, $P<0.01$; ****,$P < .0001$
